# Evaluations of short telomere risk-associated single nucleotide polymorphisms in telomerase reverse transcriptase gene on telomere length maintenance

**DOI:** 10.1101/552133

**Authors:** Jialin Xu, Matthew A. Trudeau, Andrew J. Sandford, Judy M.Y. Wong

**Affiliations:** Faculty of Pharmaceutical Sciences, University of British Columbia, Vancouver, BC, Canada; Centre for Heart Lung Innovation, University of British Columbia and St Paul’s Hospital, Vancouver, BC, Canada

## Abstract

Telomere biology disorders (TBDs) refer to a spectrum of tissue degenerative disorders caused by genetic mutations in telomere biology genes. Most patients with TBDs suffer from telomere maintenance defects secondary to telomerase deficiency. While the highly penetrant mutations in the telomerase reverse transcriptase (*TERT*) gene that drive disease onset and progression of TBDs are relatively rare, there exist several single nucleotide polymorphisms (SNPs) in *TERT* that have been linked to various diseases in the TBD spectrum. In this study, we investigated the biochemical properties of five *TERT* variants. In an *ex vivo* cell model, we found that primary human fibroblasts expressing nonsynonymous *TERT* SNPs had comparable cell growth kinetics to primary cells expressing WT-TERT, while a parallel vector control expressing-cell line entered replicative senescence. At the molecular level, primary cells expressing the minor alleles of two of the five TERT variants (A279T, ΔE441) had replication-dependent loss of telomere length. In an *in vitro* primer extension assay, these two variants showed reduced telomerase nucleotide addition processivity. Together, our data suggested that selective, common *TERT* variants could be revealed to harbour telomere maintenance defects, leading to a plausible explanation for their observed associations to telomere biology disorders.

## INTRODUCTION

Telomere biology disorders (TBDs) consist of a spectrum of tissue degenerative disorders that have variable clinical manifestations and severities, but share the same disease etiology – short telomeres [1]. Telomeres are DNA-protein structures located at the ends of eukaryotic chromosomes [2]. Telomeres shorten with cell divisions and they limit the proliferation of human cells by inducing cellular senescence when a critical length is reached. Telomerase is a ribonucleoprotein complex that can copy the template within its RNA component, telomerase RNA (TER), to telomeric DNA by using its own polymerase, telomerase reverse transcriptase (TERT). *De novo* synthesis of telomeric repeats at chromosome ends by telomerase delays the depletion of telomeres and extends the proliferation lifespan [3].

At the cellular level, mean telomere length is dynamically determined by the inherited starting telomere length, the rate of cell turnover, and the rate of telomere synthesis if telomerase is expressed in the cell. Individuals with the shortest telomeres (first percentile) in the general population are susceptible to a spectrum of premature aging disorders [1], including idiopathic pulmonary fibrosis (IPF), even if they did not carry mutations in known telomere biology genes [4].

At the organismal level, TBDs, mostly caused by genetic deficiencies in telomerase function, affect tissues’ regenerative potential and present clinically as diverse disorders. Bone marrow is one tissue that is particularly sensitive to short telomeres, as haematopoietic renewal failure occurs when the replicative potential and the stem cell pool of an affected individual is depleted. Consequently, bone marrow failure diseases, including dyskeratosis congenita (DC) and aplastic anemia, are frequently observed in patients with severe telomerase dysfunction. Since telomere shortening is essential for cell transformation and very short telomeres are found in precancerous lesions [5], patients with TBDs are at risk of developing cancers of hematopoietic and epithelial origin – i.e., cancers in high turnover tissues [6]. The lung is also frequently affected by telomere dysfunction [1, 7]. Patients with mutations in *TERT* suffer from IPF in an autosomal dominant pattern due to telomerase haploinsufficiency and the consequent short telomere-induced tissue renewal defects [8]. Furthermore, pulmonary fibrosis is found in up to 20% of all patients with DC, the prototype and the first identified telomere biology disorder [9]. Disease presentation in the lung is driven by stem cell proliferative failure following telomere dysfunction in type 2 alveolar epithelial cells (AEC2) [10].

Single nucleotide polymorphisms (SNPs) in the *TERT* region have been linked to TBDs, including, but not limited to, bone marrow failure [11, 12], aplastic anemia [12], myeloid dysplastic syndrome (a type of haematopoietic stem cell deficiency disorder) [13], combined pulmonary fibrosis and emphysema [14], and cancers of hematopoietic or epithelial origins [15–17]. While informative, these association studies did not provide a direct proof that these *TERT* SNPs’ contribution to the pathogenesis of the associated diseases. Neither did they explain how the majority of SNP carriers in the general population remain disease-free. In this study, we are interested in understanding the biological mechanisms that may explain the observed associations between common *TERT* SNPs and various TBDs. Specifically, we would like to know whether common *TERT* SNPs, when expressed in a continuous cell model over an extensive period, could modify telomerase activity and change telomere length maintenance dynamics.

## METHODS

### Cell culture

The primary human foreskin fibroblast BJ cell line was obtained from ATCC. All genetically-modified primary cell lines were cultured in DMEM high-glucose medium (Gibco, ThermoFisher Scientific, Waltham, MA, USA) with 15% fetal bovine serum (Gibco) and were maintained at 37°C with 5% CO2. Population doubling levels (PDLs) were calculated based on the following equation: PDL = (log Xe – log Xb)/0.3 + S, where Xe was the cell harvest number, Xb was the inoculum number, and S was the starting PDL.

293T is a human embryonic kidney cell transformed with adenovirus. It provides the high efficiency of transient transfection that is required for producing retrovirus packaging materials. Obtained from ATCC. 293T cell line was cultured in DMEM high-glucose medium with 5% HyClone® fetal bovine serum (GE Healthcare, Piscataway, NJ, USA) and were maintained at 37°C with 5% CO2.

### Retroviral gene delivery

The retroviral protein expression vector pBabe-hygro-Flag3B-WT-hTERT (a gift from Dr. Kathleen Collins) was used as the template for *DpnI*-mediated site-directed mutagenesis. Details for the TERT variants involved in this study are listed in S1 Table. Primers for the mutagenesis were synthesized by IDT DNA (Coralville, IA, USA) and are listed in S2 Table. The fidelity of positive clones was confirmed with Sanger sequencing of the entire TERT coding sequence (Genewiz, South Plainfield, NJ, USA). Silent mutations were introduced in each pair of primers to create a restriction site for easy identification of positive clones through restriction digestion with a selected endonuclease. The restriction enzymes of choice are also listed in S2 Table. Retroviral infection was performed by the 293T cell-based system [18] to integrate TERT (both WT and variant) coding sequences into the BJ genome. Hygromycin selection at 50 μg/ml was applied to select for positive clones for at least 14 days. Positive clones were pooled for continuous culture. Biological repeats of cell lines were performed for a selected set of *TERT* SNPs.

### Telomere length measurement

The Southern blot-based terminal restriction fragment analysis (TRF) was applied to determine average telomere lengths in cell populations [19]. Data from TRF blots were analyzed by ImageQuant Software (GE Healthcare, Piscataway, NJ, USA) using weighted average analyses [19]. Data were reported as the nearest kilobase using the 1 kb DNA ladder (New England Biolabs, Beverly, MA, USA) as a reference.

### Protein extraction

Cells were removed from culture flasks with trypsin/EDTA. Cell numbers in 100 μL of suspension were counted by Coulter Counter (Beckman Coulter, Brea, CA, USA) according to the manufacturer’s instruction and total cell numbers were calculated accordingly. Following the 2x wash with ice-cold PBS, cells were resuspended in hypotonic lysis buffer (20 mM HEPES pH 8.0, 2 mM MgCl_2_, 0.2 mM EGTA, 10% glycerol, 1 mM DTT, and 0.1 mM PMSF). Cell suspensions were subjected to four consecutive freeze-thaw cycles from liquid nitrogen to a 37 °C water bath. NaCl was then added to the lysate in two parts to a final concentration of 400 mM. Cell suspensions were incubated on ice for 15 min before clearing the lysate by centrifugation at 13,200 rpm for 15 min. The supernatant (whole cell lysate, WCL) was transferred to a fresh tube, aliquoted, and stored in −80 °C for further analysis. Protein concentrations were measured by Bradford protein assay.

### Telomerase activity

The classical radioactive primer extension assay was applied to the primary BJ cell lines expressing recombinant TERT isoforms. The quantifications of telomerase catalysis and repeat addition processivity were previously described [20].

### Immunoblot

100 μg WCL were resolved in 5% SDS-PAGE gel. A standard western blotting protocol was applied for the detection of TERT expression in the cell lysates. A sheep anti-human TERT polyclonal antibody (0.5 μg/ml, Abbexa, Cambridge, UK) and a Donkey anti-sheep horseradish peroxidase-conjugated antibody (0.2 μg/ml, Jackson ImmunoResearch, West Grove, PA, USA) were sequentially applied before detection by ECL (PerkinElmer, Waltham, MA, USA) on Kodak X-ray films. The cytoskeletal protein vinculin was used as a normalization control. It has a molecular weight of 124 kDa, which is close to TERT (127 kDa). Rabbit anti-human vinculin monoclonal antibody (0.1 μg/ml, Cell Signaling, Danvers, MA, USA) and Alexa Fluor 680 Dye (ThermoFisher Scientific, Waltham, MA, USA) were applied before detection by Licor Odyssey CLx Infrared Imaging System (LI-COR Biotechnology, Lincoln, NE, USA) and quantification by ImageJ software (NIH, Bethesda, MD, USA).

### Statistical analysis

Unpaired t-test was applied in cell lines expressing either TERT variants or WT-TERT to assess the difference between the two groups in their protein expression level, nucleotide addition processivity and repeat addition processivity. Figures were generated by GraphPad Prism 7.0 version (La Jolla, CA, US).

## RESULTS

### 1. TERT variant-expressing cells showed growth kinetics comparable to cells expressing WT-TERT

We started out by searching for common *TERT* SNPs that have been linked to diseases caused by potential short telomeres.

The nonsynonymous *TERT* SNP rs61748181 features a C>T nucleotide substitution, resulting in an amino acid change from alanine to threonine (TERT p.A279T, expressed as A279T-TERT hereafter). It is located at a linker domain of the N-terminus of TERT. Targeted deep sequencing has linked this SNP to bone marrow failure [11], lung cancer [15] and esophageal cancer [16]. Compared to other disease-associated *TERT* SNPs, it has a relatively high minor allele frequency (MAF) of 3.58% [21].

The *TERT* SNP rs34094720 features an amino acid change from histidine to tyrosine (TERT p.H412Y). Similar to the conversion from arginine to threonine in the cases of rs61748181 and rs35719940, the switch from a positively charged (basic) amino acid in rs34094720 (histidine) to an uncharged one (tyrosine) at neutral pH in the TRBD domain of TERT could potentially have profound effect on protein function. Indeed, it was first identified in two unrelated patients with aplastic anemia [12]. Later, it was also found in patients with autosomal recessive DC [22], IPF [4, 23], bone marrow failure [24], systemic sclerosis (an autoimmune connective tissue disease featured by fibrosis) [25] and in healthy volunteers [4, 24]. Despite the consistent identification of this SNP in various association studies to TBDs, there were contrasting reports about its telomerase catalytic activity: while p.H412Y was reported to have drastic reduction in telomerase catalytic activity by TRAP assay [12, 24], studies in other groups revealed a mild reduction or no change in activity by the primer extension assay [4, 26].

The *TERT* SNP rs377639087 features the deletion of codon 441 (TERT p. ∆E441). It has been discovered in patients with typical telomere biology disorders, including aplastic anemia [12], acute myeloid leukemia [27], liver cirrhosis [28], and cancer of epithelial origin such as renal cell carcinoma [29]. As this rare coding variant is known to have reduced telomerase activity and confer short telomeres [27], it served as a known telomerase activity defective control for our human cell model study. Another *TERT* SNP rs112614087 (TERT p.A615) had a heterozygosity of 0.500 [30] and therefore is deemed unlikely to change TERT function. Most of the other disease-associated *TERT* variants have relatively low MAF in Caucasians. The details of the *TERT* SNPs involved in the biochemical study are summarized in S1 Table and illustrated in the schematic of TERT functional domains in S1 Figure.

We applied an *ex vivo* human cell model to check whether the minor alleles of five TERT variants, when translated into variant versions of TERT, were functionally different from WT-TERT with long-term telomere length maintenance as a readout. We used site-directed mutagenesis to introduce the sequences encoded by the minor alleles of the SNPs into TERT open reading frame. Like other human fibroblasts, the human foreskin fibroblast BJ cell line silenced its endogenous TERT expression, whereas its TER expression is intact. Stable genomic integration of the coding sequences of TERT (WT and the SNPs) through infection with retrovirus vectors (pBabe-Hygro-flag3B-TERT) in BJ cells restored telomerase activity (Fig 1A). An examination of protein expression revealed that TERT protein levels in these reconstituted cell lines were comparable to each other (Fig 1A). The population growth kinetics of the TERT-expressing BJ cell lines followed typical logarithmic curves. While the BJ vector control cell line (BJ-vector), which does not express telomerase activity, entered replicative senescence following 10 PDLs post-antibiotic selection, all BJ fibroblasts expressing recombinant telomerase (WT/SNP variants) were proliferating continuously in our ex vivo cell model system. Notably, WT-TERT and TERT variant-expressing cell lines did not show any visible difference in growth kinetics up to 50 – 60 PDLs (Fig 1B).

**Figure 1.**
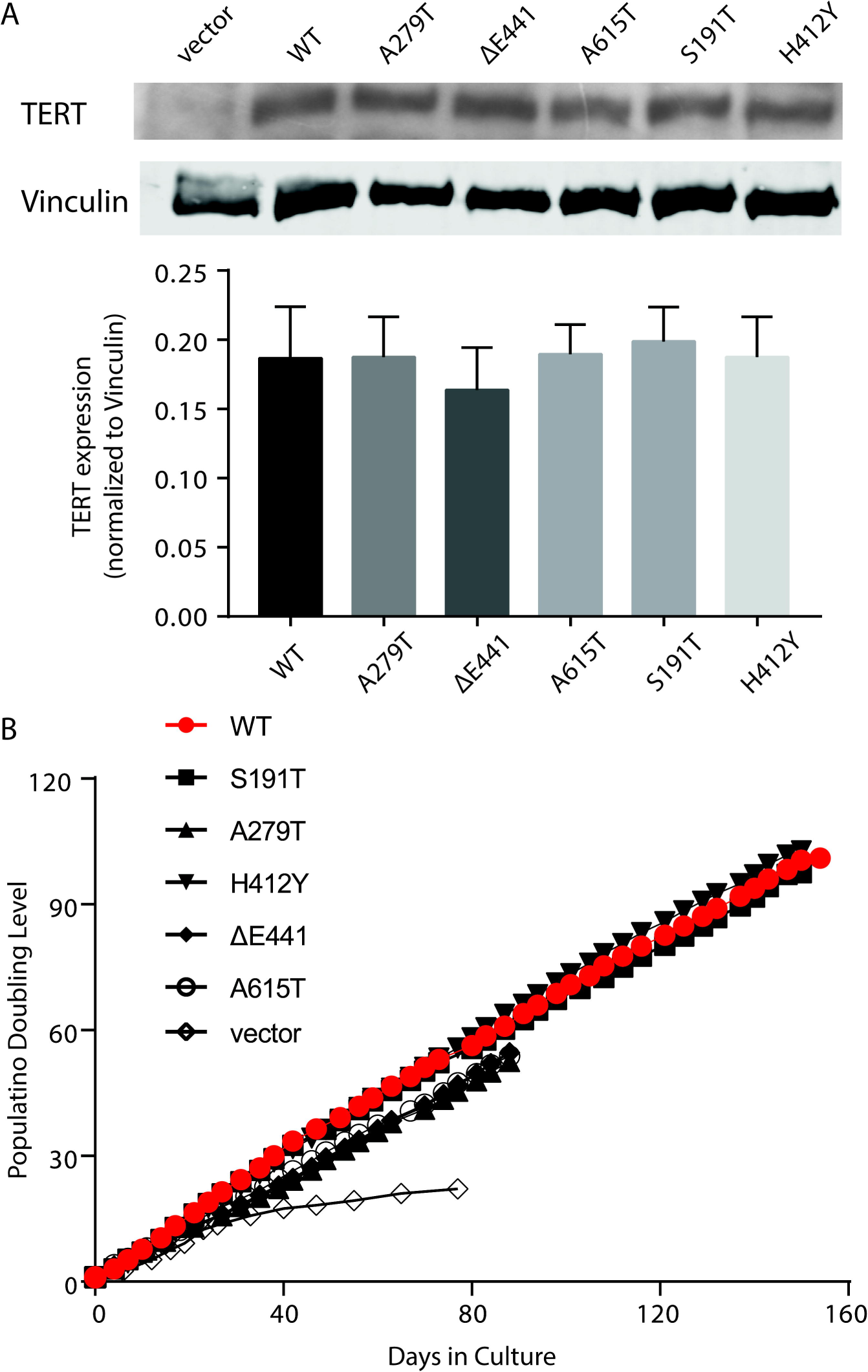
Restoration of recombinant telomerase mediated by the TERT variants in BJ cell lines. (A) Recombinant expression of TERT in BJ fibroblasts. TERT expression levels (127 kDa) are normalized to those of vinculin (124 kDa). TERT expression in different cell lines is comparable and the data are presented as mean ± SEM (n = 6). (B) Cellular growth kinetics of BJ cell lines expressing recombinant variant-version of telomerase.

### 2. Two *TERT* variants in TERT lead to compromised telomere length maintenance

Next, we checked the dynamics of mean telomere length in these cell populations over time. The measurement of telomere length confirmed that senescence in the BJ-vector cell line was caused by progressive loss of telomeric repeats (Fig 2A-C). In contrast to their comparable cellular growth kinetics, WT-BJ and variant-BJ cell lines presented differential telomere length dynamics. While BJ cell lines expressing WT-TERT and S191T, H412Y and A615T-TERT variants showed moderate telomere lengthening as a function of proliferation (Fig 2D), BJ cell lines expressing A279T and ΔE441-TERT variants had proliferation-dependent decreases in TRF within the same period (Fig 2D). In short, while the restoration of telomerase activity mediated by some *TERT* SNPs (i.e., A279T and ΔE441) prevented BJ cells from entering replicative senescence, these cell lines showed compromised telomere maintenance capacity within 40 PDLs, compared to WT-TERT expressing cells.

**Figure 2.**
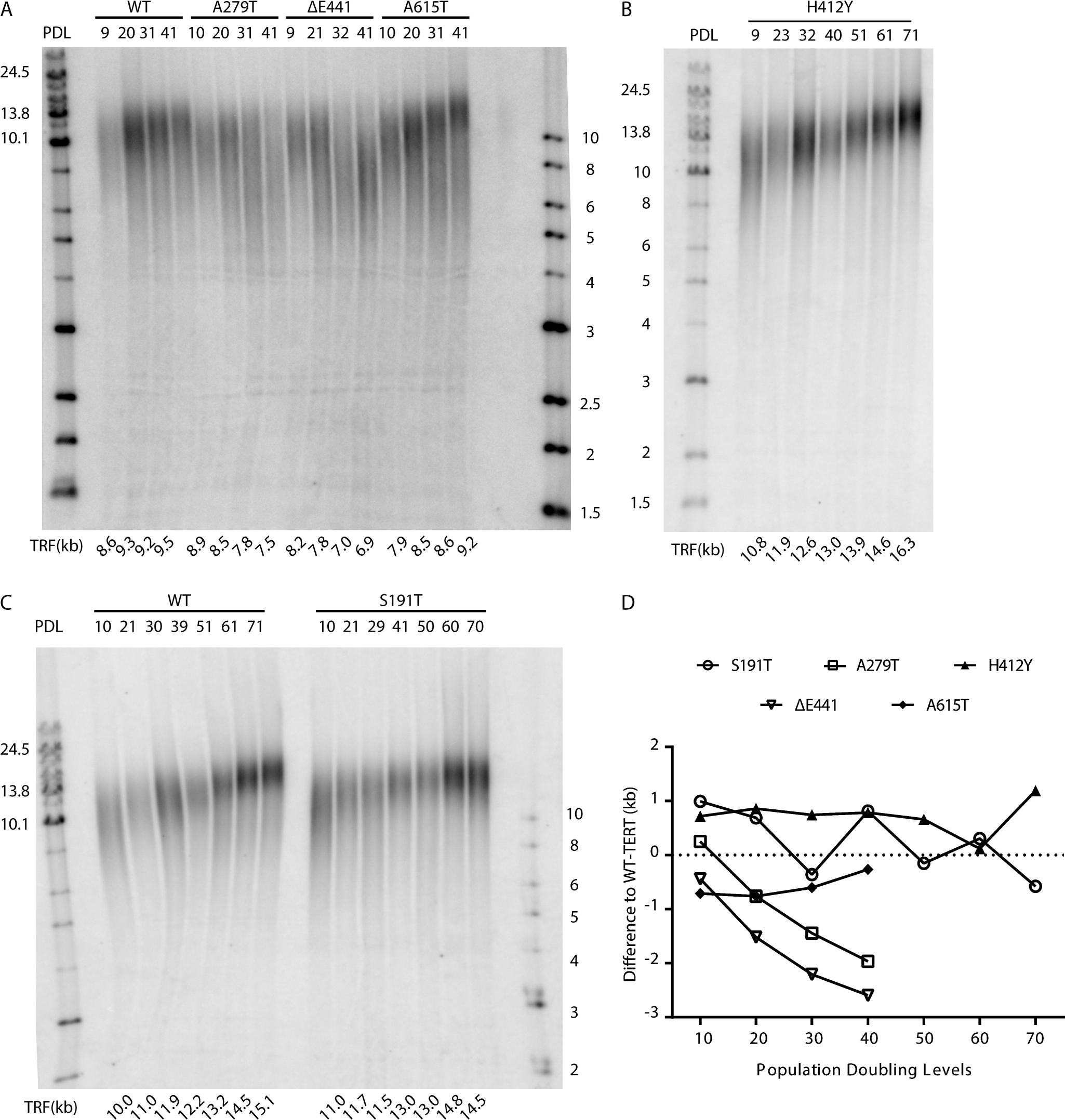
Defective telomere length maintenance was found in cells expressing A279T and ΔE441. (A-C) Terminal restriction fragment analysis was used to quantify telomere lengths. The measured signal represents the sum of telomeric and sub-telomeric region. Estimated telomere length was calculated by the weighted average of the lane and is shown at the bottom of the gel. All samples tested here were collected from cell populations with different doubling levels. The PDL for each lane is also shown on the top of the gel. (D) Two out of five TERT variants (A279T and ΔE441) conferred telomere length maintenance defects compared to WT-TERT. The other three TERT variants (S191T, H412Y and A615T) showed comparable telomere length increment to WT.

### 3. Defective telomere length maintenance of *TERT* variants is caused by reduced telomerase activity and processivity

To understand the mechanism behind defective telomere length maintenance by TERT variants, we measured telomerase activity *in vitro* for the cell lines expressing A279T and ΔE441-TERT.

We applied the primer extension assay to check single nucleotide incorporation (nucleotide addition processivity, NAP) and repeat synthesis (repeat addition processivity, RAP) using cell extracts from WT-TERT and TERT variant cell lines. In this assay, an 18 nt telomeric oligonucleotide primer ending with TTA is extended in the presence of [α-^32^P]-dGTP. With this specific primer, telomerase incorporates a single radiolabeled guanidine, stalls and translocates as it reaches the end of the TER template region. As a result, the characteristic 6 nt telomerase repeat pattern is observed, starting at TTA-primer +1, +7, +13, and so on (Fig 3A). Within each repeat, the guanosines reside in TTA-primer +5 and +6 positions are also radiolabeled at the α phosphate position, yet these guanosines are in the middle of the templated repeat and the chance of telomerase stalling at these locations is low. As such, they do not show signals as strong as guanosines at the TTA-primer +1 do. The activities are normalized to a radioactive end-labeled 15 nt oligo recovery control so that the DNA extraction/purification efficiency in different samples are comparable.

**Figure 3.**
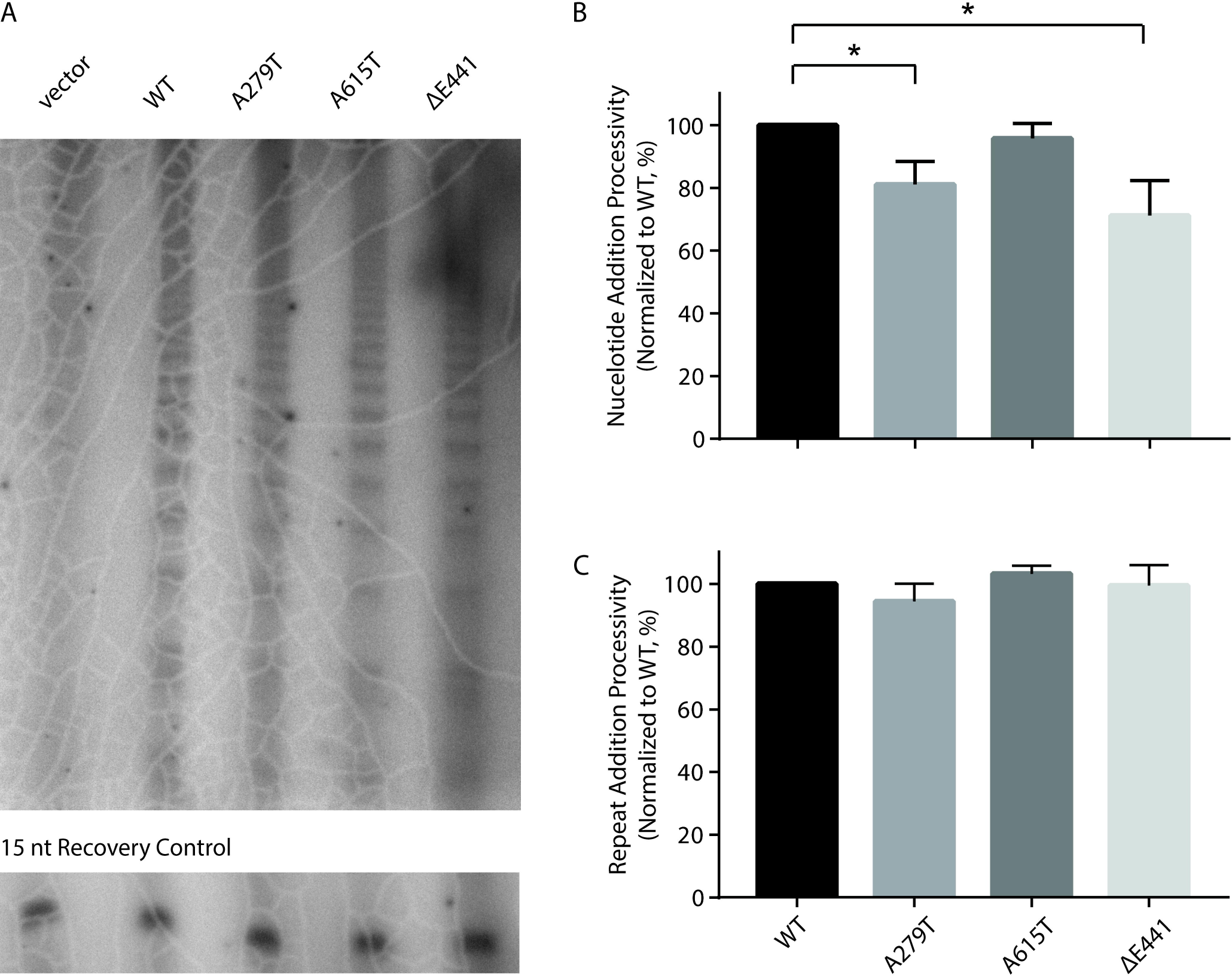
*TERT* variants affect telomerase activity and processivity in a SNP-specific manner. (A) Expression of variant TERT proteins in BJ cell lines in the primer extension assay. RC, recovery control. (B) Nucleotide addition processivity in BJ cell lines expressing TERT variants. (C) Repeat addition processivity in BJ cell lines expressing TERT variants. Data are presented as mean ± SEM (n=6).

The NAP and RAP activities mediated by the TERT variants were normalized to WT and illustrated in Fig 3B & 3C and summarized in Table 1. The telomerase from all the genetically-modified BJ cell lines under investigation showed comparable RAP. However, telomerase from three of the five cell lines (BJ cell lines expressing A279T, ΔE441, and S948R) showed slight but statistically significant decreases in their NAP activities. For example, the RAP activity of the A279T-TERT was 94.4 ± 5.5% compared to the WT-TERT (n = 6, p = 0.3313). Its NAP activity was 81.1 ± 7.3% of WT (n = 6, p = 0.0323). The overall trend showed that the TERT protein encoded by A279T had similar repeat addition capacity as WT-TERT, but reduced nucleotide addition processivity.

**Table 1.** Summary of NAP and RAP activity for the selected *TERT* SNPs

| Variant | Nucleotide Addition Processivity<sup>\$</sup> | | | Repeat Addition Processivity<sup>\$</sup> | | |
| --- | --- | --- | --- | --- | --- | --- |
|  | Mean | SEM | *P* Value | Mean | SEM | *P* Value |
| A279T | 81.05% | 7.33% | 0.032<sup>*</sup> | 94.41% | 5.52% | 0.331 |
| ΔE441 | 71.16% | 11.22% | 0.022<sup>*</sup> | 99.41% | 6.48% | 0.929 |
| A615T | 95.78% | 4.77% | 0.347 | 103.2% | 2.50% | 0.110 |

<sup>\*</sup> Both NAP and RAP are normalized to WT-TERT. These data are calculated based on six biological repeats.
<sup>\$</sup> $p < 0.05$ .

## DISCUSSION

We investigated the biological effects of variants in the *TERT* gene that have previously been linked to telomere biology disorders. Using a combination of molecular and cell biology tools in the laboratory, we have demonstrated that two out of five nonsynonymous *TERT* variants could lead to deficient telomere length maintenance.

Our *ex vivo* results demonstrated in a pure laboratory setting that, when other confounding factors for telomere attrition were fixed, fibroblasts with recombinant expressions of several disease-associating TERT variants were predetermined to suboptimal telomere maintenance. However, deficient telomere length maintenance in this context did not translate to a loss of proliferative capacity, as BJ cells expressing the minor alleles of the TERT variants were kept in culture for up to 50 – 60 PDLs, the limit of fibroblast divisions during a lifetime [31], and demonstrated the same growth rate as WT-TERT expressing cells. To our knowledge, this was the first study to investigate the long-term effects of common TERT variants on telomeres in primary cell lines. Our results confirmed that the short telomere risk-associated SNPs encoded TERT variants A279T and ΔE441 that were deficient in telomere length maintenance. It provided experimental evidence to support the previous clinical observations that in subjects with bone marrow failure, carriers (heterozygous and homozygous) of these variants had shorter telomeres than their age-matched peers [11]. Our investigation of their biochemical properties revealed that while the variants’ RAP activities were intact, both A279T and ΔE441 displayed slightly lower NAP activity compared to WT-TERT. These biochemical results are also supported by reports in different cellular and experimental systems [16, 26].

Interestingly, in cell model over long-term culture (up to 70 PDLs), we also revealed that the short telomere risk-associated TERT variant H412Y had telomere length maintenance capacity comparable to WT-TERT. Hence, its frequent identification in association studies to various TBDs does not indicate that carriers of this variants have short telomeres. While H412Y itself does not cause short telomeres, it remains possible that this rare variant is in linkage disequilibrium with another yet unidentified variant that is important in telomere length maintenance that contributed to the pathogenesis of TBDs.

An individual’s telomere length is determined by multiple factors, including genetics, inherited telomere length, and environmental factors. While the effects of environmental insults, such as smoking or infection in the case of lung diseases, were subject to huge inter-individual and temporal variations, the effects represented by the inherited telomere length and from the genotype of telomere-related genes were fixed. The comprehension of these fixed effects could facilitate understanding of the pathogenesis and progression in complex disorders.

In this study, we revealed recombinant expression of common TERT variants in a primary human cell line exhibited replication-dependent loss of telomere lengths. The minor defects in telomere length maintenance did not translate into cellular growth defects within the duration of our experiment (60 - 100 PDLs). Accordingly, carriers of these common genetic variants, in the absence of additional factors (environmental or genetic) that accelerate telomere attritions, are not expected to display pathogenesis associated with short telomeres. However, carriers of (even a single) defective common variant in telomere biology genes are still genetically predetermined to a decreased capacity to maintain their telomeres, when compared with noncarriers. These deficiencies, while silent phenotypically within the general population, could be exacerbating the clinical presentations of tissue degenerative conditions, as seen in the increase in cell renewal demands from the deleterious effects of smoking or recurrent infections. Under these constant renewal pressures, the minor defects in telomerase catalysis in the variant carriers could be revealed as a clinical determinant. While these minor defects in telomerase could not lead to monogenic disorders, we contend that they do contribute to the sum total of all etiologic factors (genetic and environment) leading to the loss of tissue integrity and perhaps earlier onset as well as faster progression of many aging-related diseases, such as cancer [33], idiopathic pulmonary fibrosis [34], or COPD/emphysema [34]. With this hypothesis, carriers of common defective variants in telomerase and telomere biology genes should experience accelerated decline of existing tissue degenerative conditions. This should provide an impetus for novel epidemiological and clinical evaluations of these common variants on their association with the progression rather than the risk of a tissue-degenerative disorder.

## CONCLUSIONS

In summary, we investigated telomere length maintenance effect for a panel of variants in *TERT* in *ex vivo* cellular models. While three TERT variants (S191T, H412Y, and A615T) showed comparable telomere maintenance dynamics to WT-TERT, two TERT variants (A279T and ΔE441) showed defective telomere maintenance secondary to reduced telomerase nucleotide addition processivity.

## Supporting Information Captions

**S1 Table.** List of *TERT* SNPs: nucleotide and amino acid changes and their frequencies

**S2 Table.** Primer design for site-directed mutagenesis

**S1 Figure.** Schematic of TERT functional domains, showing the variants involved in this study.

Nonsynonymous amino acid changes in this study are illustrated in the schematic of TERT functional domains. Studies from orthologs in other species, and from biochemical studies of human telomerase reverse transcriptase have mapped the following functional features in the TERT protein. It has a telomerase essential N-terminal domain (TEN) (1 – 195 aa), the telomerase RNA-binding domain (TRBD) (322 – 594 aa), the reverse transcriptase (RT) domain (595 – 935 aa), and the C-terminal extension (CTE) (936 – 1132 aa).

