## Supplementary Tables and Figure for "Evaluations of short telomere risk-associated single nucleotide polymorphisms in telomerase reverse transcriptase gene on telomere length maintenance"

**S1 Table.** List of *TERT* SNPs: nucleotide and amino acid changes and their frequencies

| Amino Acid Change | Nucleotide Change | dbSNP Number | Overall | | Caucasians | | MAF | | Functional domain[$] | Data source[#] |
|---|---|---|---|---|---|---|---|---|---|---|
| | | | | | | | Overall | Caucasians | | |
| S191T | 572G>C | rs11952056 | 2 | 5006 | 0 | 1006 | 0.040% | 0.000% | TEN | [1] |
| A279T | 835G>A | rs61748181 | 48 | 4960 | 36 | 970 | 0.958% | 3.579% | between TEN and TRBD | [1] |
| H412Y | 1234C>T | rs34094720 | 3 | 5005 | 3 | 1003 | 0.060% | 0.298% | TRBD | [1] |
| E441del | 1323_1325 delGGA | rs377639087 | 4 | 5004 | 4 | 1002 | 0.080% | 0.398% | TRBD | [1] |
| A615T | 1843G>A | rs112614087 | 3 | 246159 | 1 | 111629 | 0.001%[*] | 0.001% | RT | [2] |

[#] Data retrieval date: August 31, 2018.

[*] The heterozygosity of rs112614087 was listed as 0.500 in the NCBI dbSNP database when we initiated the study in 2013 [3]. Therefore, during the initiation of this study, this variant was deemed unlikely to change TERT function and was selected as a known telomerase activity intact control for our human cell model study.

[$] TEN, telomerase essential N-terminal domain; TRBD, telomerase RNA binding domain; RT, reverse transcriptase domain.

**S2 Table.** Primer design for site-directed mutagenesis

| Primer # | dbSNP ID | | Sequence (5' – 3') |
|---|---|---|---|
| TERT S191T with *Age*I-F | rs11952056 | Forward | CCC CGC CAC ACG CTA CCG GTC CCC GAA GGC GT |
| TERT S191T with *Age*I -R | | Reverse | ACG CCT TCG GGG ACC GGT AGC GTG TGG CGG GG |
| TERT A279T with *Eag*I-F | rs61748181 | Forward | GTG GTG TCA CCT GCC CGG CCG ACC GAA GAA GCC ACC TCT |
| TERT A279T with *Eag*I-R | | Reverse | AGA GGT GGC TTC TTC GGT CGG CCG GGC AGG TGA CAC CAC |
| TERT H412Y with *Bbs*I-F | rs34094720 | Forward | CCC TAC GGG GTG CTC TTG AAG ACG TAC TGC CCG CTG CGA |
| TERT H412Y with *Bbs*I-R | | Reverse | TCG CAG CGG GCA GTA CGT CTT CAA GAG CAC CCC GTA GGG |
| TERT E441del with *Bbs*I-F | rs377639087 | Forward | GGC GGC CCC CGA GGA AGA CAC AGA CCC CCG T |
| TERT E441del with *Bbs*I-R | | Reverse | ACG GGG GTC TGT GTC TTC CTC GGG GGC CGC C |
| TERT A615T-*Hinc*II-F | rs112614087 | Forward | GAA GCC AGG CCC ACC CTG TTG ACG TCC AGA C |
| TERT A615T-*Hinc*II-R | | Reverse | GTC TGG ACG TCA ACA GGG TGG GCC TGG CTT C |

**S1 Figure.** Schematic of TERT functional domains, showing the variants in this study.

Nonsynonymous amino acid changes in this study are illustrated in the schematic of TERT functional domains. Studies from orthologs in other species, and from biochemical studies of human telomerase reverse transcriptase have mapped the following functional features in the TERT protein. It has a telomerase essential N-terminal domain (TEN) (1 – 195 aa), the telomerase RNA-binding domain (TRBD) (322 – 594 aa), the reverse transcriptase (RT) domain (595 – 935 aa), and the C-terminal extension (CTE) (936 – 1132 aa).

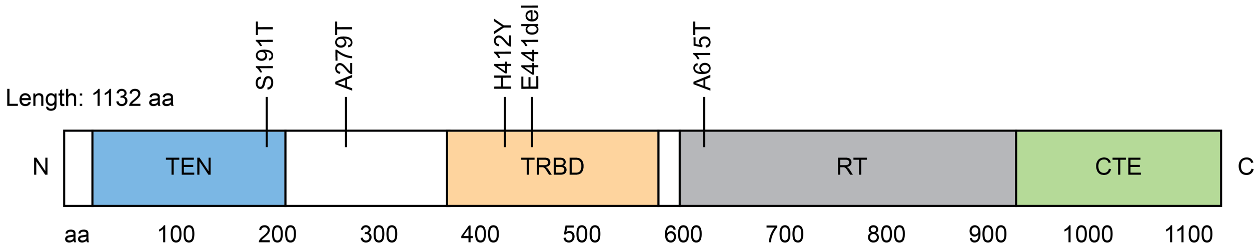


1. Auton A, Brooks LD, Durbin RM, Garrison EP, Kang HM, Korbel JO, et al. A global reference for human genetic variation. Nature. 2015;526(7571):68-74. doi: 10.1038/nature15393. PubMed PMID: 26432245; PubMed Central PMCID: PMCPMC4750478.

2. Lek M, Karczewski KJ, Minikel EV, Samocha KE, Banks E, Fennell T, et al. Analysis of protein-coding genetic variation in 60,706 humans. Nature. 2016;536(7616):285-91. doi: 10.1038/nature19057. PubMed PMID: 27535533; PubMed Central PMCID: PMCPMC5018207.

3. Sherry ST, Ward MH, Kholodov M, Baker J, Phan L, Smigielski EM, et al. dbSNP: the NCBI database of genetic variation. Nucleic Acids Res. 2001;29(1):308-11. PubMed PMID: 11125122; PubMed Central PMCID: PMCPMC29783.
